# Exploring Neural Dynamics in Self-Voice Processing and Perception: Implications for Hallucination Proneness

**DOI:** 10.1101/2023.09.21.558843

**Authors:** Suvarnalata Xanthate Duggirala, Hanna Honcamp, Michael Schwartze, Therese van Amelsvoort, Ana P. Pinheiro, David E. J. Linden, Sonja A. Kotz

## Abstract

Altered sensory feedback processing and attention control are assumed to contribute to auditory verbal hallucinations, which are experienced by the general population and patients with psychosis, implying a continuum of hallucination proneness (HP). However, the interaction of altered sensory feedback processing and attention control along this HP continuum remains unclear. Manipulating the level of certainty of sensory feedback by changing self-voice quality (100% neutral, 60-40% neutral-angry, 50-50% neutral-angry, 40-60% neutral-angry, 100% angry) in individuals varying in HP, we tested this interaction using electroencephalography while participants self-generated or passively listened to their voices. Regardless of voice quality, HP modulated the N100 and P200 suppression effects. High HP individuals showed an increased N100 response to the self-generated voices and an increased P200 response for externally-generated voices. This may indicate increased error awareness and attention allocation in high HP individuals for self-voice generation stemming from altered sensory feedback processing, and/or attentional control. The current findings suggest that alterations of the sensory feedback processing in self-voice production are a fundamental characteristic of the continuum of HP, regardless of the clinical status of voice hearers.

**Highlights:**

- Altered N100 voice suppression in high HP, regardless of the clinical status.
- High HP associated with altered sensory feedback processing and attentional control.
- Current findings support a ‘neurophysiological’ continuum of HP.

## 1. Introduction

Auditory verbal hallucinations (AVH) and hearing voices are perceptions without corresponding sensory input. Different cognitive models and theories have been proposed to account for AVH [1–10]. Supported by the ‘forward model’ framework, source monitoring and inner speech models of AVH postulate that voice hearers misidentify an internally generated action (i.e., speech/voice) as originating from an external source [11–15]. However, an efficient perceptual system needs to clearly differentiate internally-from externally-generated sensory events. The forward model postulates that a top-down prediction (efference copy) of the sensory consequences of an internally-generated event is produced, which is then compared to the perceived sensory feedback. Based on the degree of mismatch (prediction error) between predicted and perceived sensory feedback, the internal model is updated and the event is classified as either internally- or externally-generated. This theory is supported by evidence from the visual [16–18], tactile [14, 19, 20], and auditory domain [21–25].

Electroencephalographic (EEG) studies identified the N100 auditory evoked potential as an indicator of the degree of mismatch between predicted and perceived sensory feedback [26–28]. The N100 primarily originates in the auditory cortex (AC) [29–32] and its amplitude is suppressed when predicted and perceived sensory feedback to a self-generated voice matches closely [26–28]. Conversely, a larger degree of mismatch leads to an increased N100 response and potentially also leads to increased attention allocation [33, 34]. Altered self-other voice discrimination has been reported in both psychotic as well as non-clinical voice hearers [6, 35–37], which is evidenced by reduced N100 suppression and increased neural activity in AC for self-compared with externally-generated voice [26–28, 38–41]. This indicates that self-generated sensory events might be processed incorrectly, and might be misattributed to an external source in voice hearers regardless of their need for clinical care. Similarly, the aberrant salience hypothesis puts forward that voice hearers misattribute salience to an irrelevant stimulus because they cannot inhibit attention to them [42–44]. In empirical studies with voice hearers, misattribution of salience by allocating attention to irrelevant stimuli manifests as misattributing negative value to neutral stimuli [45] and as perceiving meaningful speech in noise [46]. Altered self-monitoring and salience misattribution theories might therefore share a common denominator through predictions [47–49]. Consequently, the lack of disengagement from prior expectations might result in increased attention allocation and attachment of meaning to non-relevant stimuli. Alterations of sensory feedback processing and attentional control therefore seem to present as two sides of the same coin in voice hearing.

The underlying neural mechanisms seem to be generic to AVH rather than psychosis-specific [50–53]. Psychotic, clinically at-risk of psychosis, and non-clinical AVH may accordingly lie on a severity continuum that ranges from low to high hallucination proneness (HP) [54–56]. This suggests that the alterations in sensory feedback processing and attentional control evidenced in psychotic [26–28] and non-clinical [38, 39] voice hearers should also be present in non-voice hearers with high HP, albeit in an attenuated form. Emotional voice quality can additionally affect sensory feedback processing and modulate AC activity [57]. The differences between non-clinical voice hearers and voice hearers with a psychotic disorder pertain to the emotional quality of their voices, controllability, and related distress [51, 58]. Psychotic voice hearers often perceive derogatory voices that are beyond their control, which causes distress and significantly impacts their daily life [51, 58]. In contrast, non-clinical voice hearers tend to perceive neutral or positive voices, can exert control over them, and rarely experience distress [51, 58]. These differences between psychotic and non-clinical voice hearers influence how they evaluate and judge negative emotional content [59–61]. Contrary to voice hearers with psychotic disorder, non-clinical voice hearers can downregulate negative emotions by dampening their emotional salience [62]. A systematic manipulation of negative emotional voice quality could thus provide insight into individuals’ certainty about sensory feedback in their own voice, highlighting a potential trade-off with attentional control in persons who vary in HP. In turn, this may be key to a better understanding of the risk of transitioning from high HP to pathological voice hearing.

The current EEG study therefore systematically investigated effects of modulated sensory feedback to one’s own voice varying in its degree of emotional quality and certainty of sensory feedback to self-voice as a function of HP. Using a well-validated auditory-motor task, participants self-generated and passively listened to their self-voice, which changed from fully neutral to fully angry: 100% neutral, 60-40% neutral-angry; 50-50% neutral-angry; 40-60% neutral-angry and 100% angry. The high temporal resolution of EEG allowed analysis of short-, mid- and long-latency event-related potential (ERP) markers to differentiate early and later auditory processing stages. Specifically, we probed sensory feedback processing and error awareness/attention allocation using the earlier N100 as primary outcome measure and then focused on the later categorical distinction of the self-from the externally-generated voice using the P200 as secondary outcome. The main hypothesis was that high HP would be linked to alterations of sensory feedback processing of the self-voice and of attentional control [39]. High HP individuals were expected to display increased N100 and P200 responses to self-as compared to externally-generated unambiguous self-voice (100% neutral and 100% angry). The self-generated ambiguous self-voice (60-40%: neutral-angry; 50-50%: neutral-angry; 40-60%: neutral-angry) was expected to result in increased N100 and P200 responses in low but not high HP individuals.

## 2. Methods

### Participants

45 adult participants, including 2 voice hearers with a diagnosis of a psychotic disorder were recruited (see supplementary document for power calculations). All participants followed the same procedure and three testing sessions. Prior to any assessment, participants were informed about the study procedures via an information letter as well as in person. Five participants could not participate in further sessions. Therefore, the final sample included 40 participants (26 females, 13 males, 1 other; mean age = 24.45, s. d = 2.33 years; range = 20-44 years) varying in HP as measured by the Launay Slade Hallucination Scale (LSHS [54, 63]) (LSHS total scores: mean = 16.55, s.d. = 12.90, max = 57, min = 0; LSHS AVH scores [sum of items: “In the past, I have had the experience of hearing a person’s voice and then found no one was there”, “I often hear a voice speaking my thoughts aloud”, and “I have been troubled by voices in my head”]: mean = 2.52, s.d. = 3.14, min = 0, max = 11). All participants provided their written informed consent. They either received financial compensation in vouchers (10 euros per hour) or study credits (1 credit per hour) for their participation. All participants self-reported normal or corrected-to-normal visual acuity and normal hearing. The study was approved by the Medical Ethics Review Committee of the azM and Maastricht University (METC azM/UM) and conducted in accordance with the Declaration of Helsinki (METC 20-035; the study was prematurely terminated due to recruitment difficulties concerning voice hearers).

### Procedure

All participants went through three testing sessions.

#### Session 1: Screening and neuropsychological assessment

The first session consisted of a diagnostic interview (the Mini International Neuropsychiatric Interview)[64] and a detailed neuropsychological assessment of voice-hearing (including LSHS). This was done to differentiate non-voice hearers, non-clinical voice hearers, and voice hearers with a psychotic disorder.

#### Session 2: Voice recording and stimulus generation

Participants comfortably sat inside an acoustically isolated chamber with the recording equipment (SENNHEISER K6/ME64 condenser microphone coupled with the M-track Eight as the audio interface), while the researcher sat outside this chamber. Recordings were made using a microphone with the Praat software (https://www.praat.org.). Participants were instructed to repeatedly vocalize “ah” in a neutral (i.e., without emotion) and in an angry voice for approximately 500 ms. They were provided with examples to familiarize them with the target duration of the vocalization. This specific duration was chosen to properly capture the emotionality while maintaining self-voice recognition. A vowel was chosen (instead of words) to eliminate semantic content processing [23, 65, 66]. The best voice sample was selected if the participants confirmed that (i) they recognized their recorded voice, (ii) the anger intensity was the highest that they could produce, (iii) they perceived no emotion in the neutral recording, and (iv) if the vocalization was pronounced clearly. Background noise was eliminated from the recordings using Audacity software (https://audacityteam.org/), and a Praat script was applied to normalize the intensity at 70 dB. The duration of the final neutral and angry “ah” vocalizations for each participant was 500 ms.

### Morphing

To create voice samples with varying degrees of emotional content, the pre-recorded neutral and angry self-voices for each individual participant were parametrically morphed to create a neutral-to-angry continuum. These continua consisted of 11 stimuli with 10% stepwise increase (neutral-to-angry) in emotional content along the continuum (supplementary table 1). Morphing was performed using TANDEM-STRAIGHT software [67–69] running on MATLAB (R2019a, v9.6.0.1072779, The MathWorks, Inc., Natick, MA). For the final EEG experiments, 100% neutral, 60-40%: neutral-angry; 50-50%: neutral-angry; 40-60%: neutral-angry and 100% angry voice morphs were selected. The intermediate voice morphs were selected based on pilot data revealing that a maximum of uncertainty to differentiate the neutral from a somewhat angry voice fell in the range of 35-65% morphing. The increase in emotional voice quality (as self-voice changes from fully neutral to fully angry) and manipulations of certainty (as the self-voice quality changes from most certain to somewhat ambiguous to certain again; most certain: 100% neutral, uncertain/ambiguous: 60-40%: neutral-angry; 50-50%: neutral-angry; 40-60%: neutral-angry; certain: 100% angry self-voice morphs) allowed probing changes in certainty of sensory voice feedback and attentional control resulting from these changes.

#### Session 3: EEG recordings

The third session comprised the EEG recordings. Participants were given an overview of the procedure and the principles of EEG at the start of the session. A previously employed auditory-motor task was used to investigate differences in self- and externally-generated auditory stimuli [39] (figure 1). Previous studies have consistently shown an N100 amplitude suppression for sounds generated by a button-press, in contrast to those generated externally [70–74], meaning that button-presses can be reliably used as a motor-act to self-generate a stimulus.

**Figure 1:**
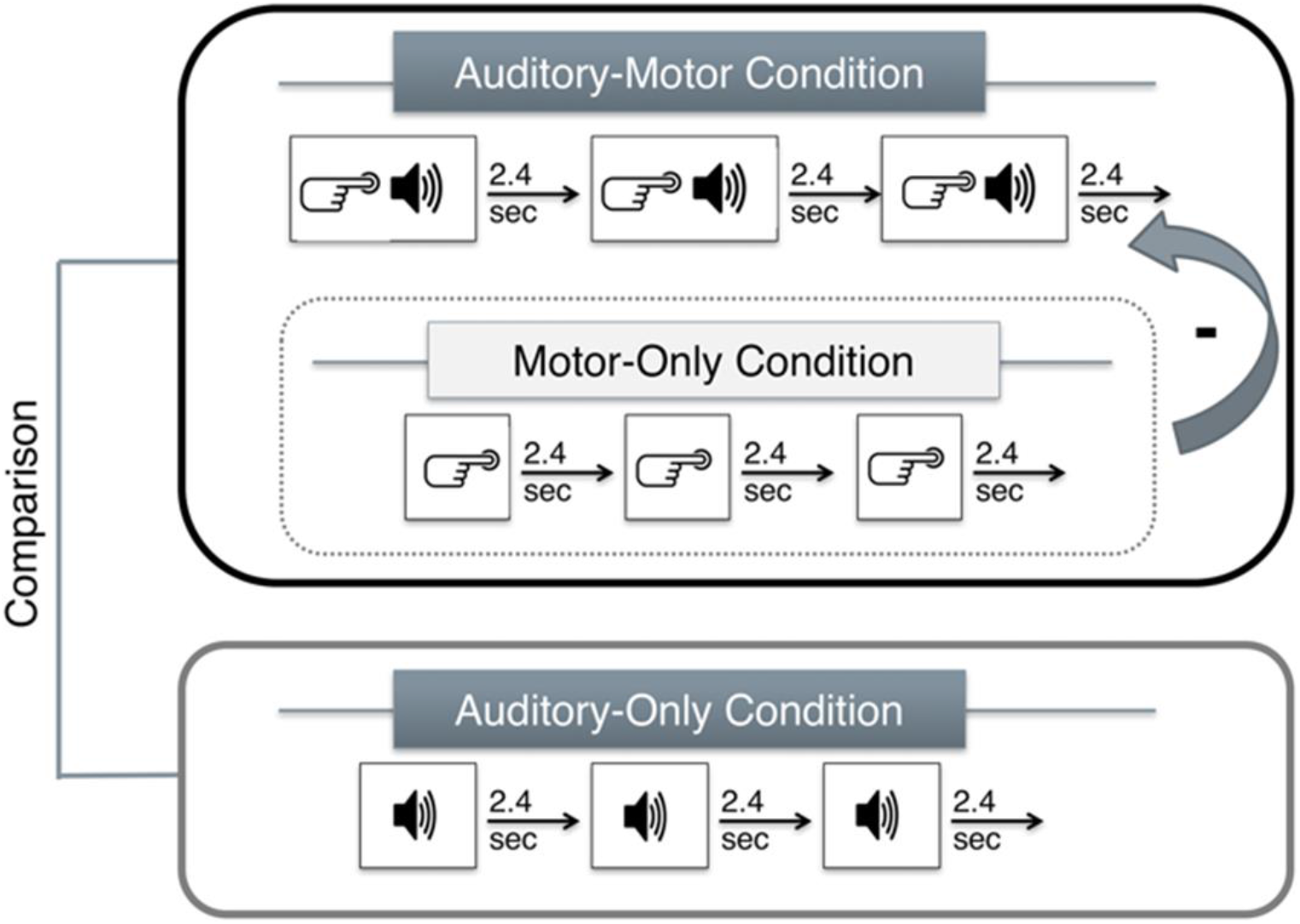
Motor-Auditory Task (Pinheiro et al., 2018). The task included three conditions: a motor-auditory (MA) condition, where participants pressed a button to generate a pre-recorded auditory stimulus representing a self-generated stimulus; an auditory-only (AO) condition, where participants passively listened to the pre-recorded auditory stimulus representing an externally-generated stimulus; and a motor-only (MO) condition, wherein they pressed a button but did not hear anything. This last condition was used to control motor activity resulting from the button-press in the MA condition (MA-MO = MA corrected [MAc]).

The paradigm was presented in a mixed design and consisted of 5 runs in total (supplementary figure 1). All conditions (MA, AO, and MO) were blocked and 5 types of stimuli were randomly presented within the MA and AO blocks. Four of these 5 runs consisted of 2 sections corresponding to MA, followed by AO. Each of these sections consisted of 10 blocks corresponding to right and left hand button presses. Each block consisted of 10 trials. Five types of voice morphs belonging to “ah” vocalizations, respectively, were presented during MA and AO conditions. Overall, this resulted in 80 trials for each stimulus type. Trial durations from the MA section were used to present the voices in the AO section within each run. Therefore, each trial duration varied according to the button press. Prior to the experimental runs, participants were trained to press the button approximately every 2.4 seconds for the MA condition consisting of 100 trials. They received feedback during the training to adjust their tapping speed. Training concluded only if they performed correct taps in 75% of the trials [39]. No feedback was provided during the experimental blocks. Participants could take small breaks after each run. The task was programmed and presented using the Presentation software (version 18.3; Neurobehavioral Systems, Inc.). Stimuli were presented via in-ear inserts. Button presses were recorded via the spacebar button on the keyboard.

At the end of the EEG session, participants were asked to rate the auditory stimuli for arousal and valence (supplementary figure 2). In addition, they were asked to rate the stimuli on perceived ownness, meaning perceived certainty of their own voice. This was done to ensure that they recognized their own voices and perceived the respective emotions.

### EEG data acquisition and processing

EEG data were recorded in an acoustically and electrically shielded room with BrainVision Recorder (Brain Products, Munich, Germany) using an ActiChamp 128-channel active electrode set-up while participants performed the task. Data were acquired at a sampling frequency of 1000Hz, electrode impedance was kept below 10 kΩ, and with FCz as online reference. During the EEG recording, participants were seated in a comfortable chair about 100 cm away from a screen in front of them.

EEG data were pre-processed using the MATLAB 2019a based toolbox Letswave 6 (https://github.com/NOCIONS/letswave6). Data were first cleaned to remove any repeated blocks/runs of the EEG recordings (e.g., due to hardware issues with the ear inserts leading to noise in the auditory stimuli or complete absence of audition), and trials within the time range of 800-2400 ms were processed further. Data were then downsampled to 500 Hz and bandpass filtered (0.5-30 Hz). All channels were re-referenced to the average of the four (TP7, TP8, TP9, TP10) mastoid electrodes. Noise related to eye blinks and movements, and noisy electrodes were removed using independent component analysis (ICA) with the runica algorithm in combination with principal component analysis (PICA; Rajan and Rayner as implemented in letswave 6). ICs representing the above mentioned noise were removed for each participant based on the IC time course and topography. The resulting data were segmented using a time-window of -400 to 800 ms relative to the onset of the auditory stimuli. The segmented data were baseline corrected to a window of -200 to -0 ms relative to the onset of the auditory stimuli. After baseline correction, an automatic artifact rejection algorithm was applied with an amplitude criterion of ± 65µV to remove epochs/trials with remaining artefacts. The resulting data were then averaged for each participant, for each condition. The grand average waveforms revealed four distinct ERP components (figure 2), two negative ones peaking at approximately 110 ms and 450 ms respectively and two positive ones peaking at 55 ms and 200 ms approximately. Mean amplitudes (P50: 0.035-0.075, range = 40 ms, N100: 0.08-0.14, range = 60 ms; P200: 0.16-0.24, range = 80 ms; N200: 0.39-0.49, range = 100 ms) extracted from one fronto-central region of interest [73, 75–78] ROI with 20 electrodes: AFF1h, AFF2h, F1, Fz, F2, FFC3h, FFC1h, FFC2h, FFC4h, FC3, FC1, FC2, FC4, FCC3h, FCC1h, FCC2h, FCC4h, C1, Cz, C2) were chosen as the outcome measure (figure 2).

**Figure 2:**
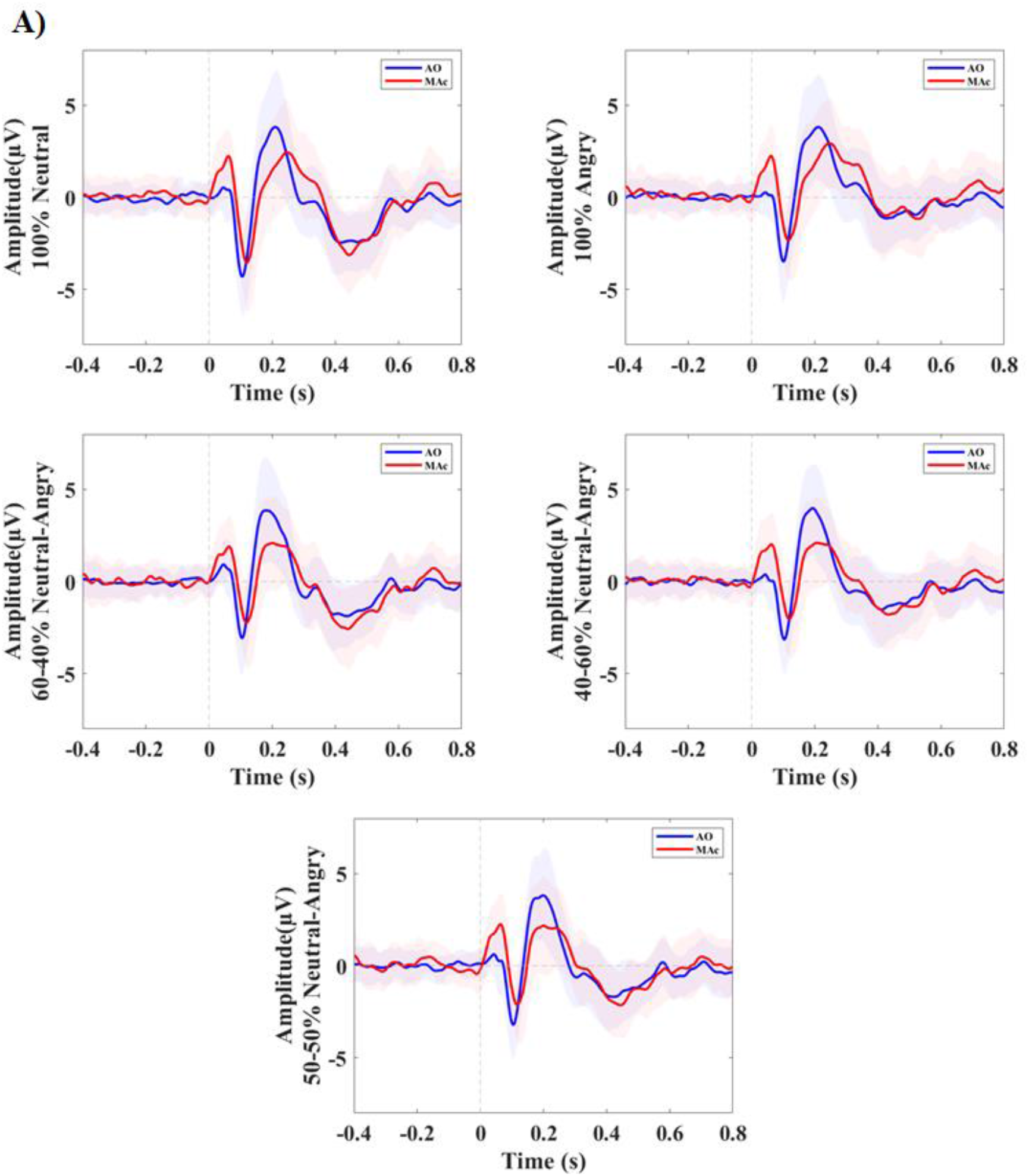
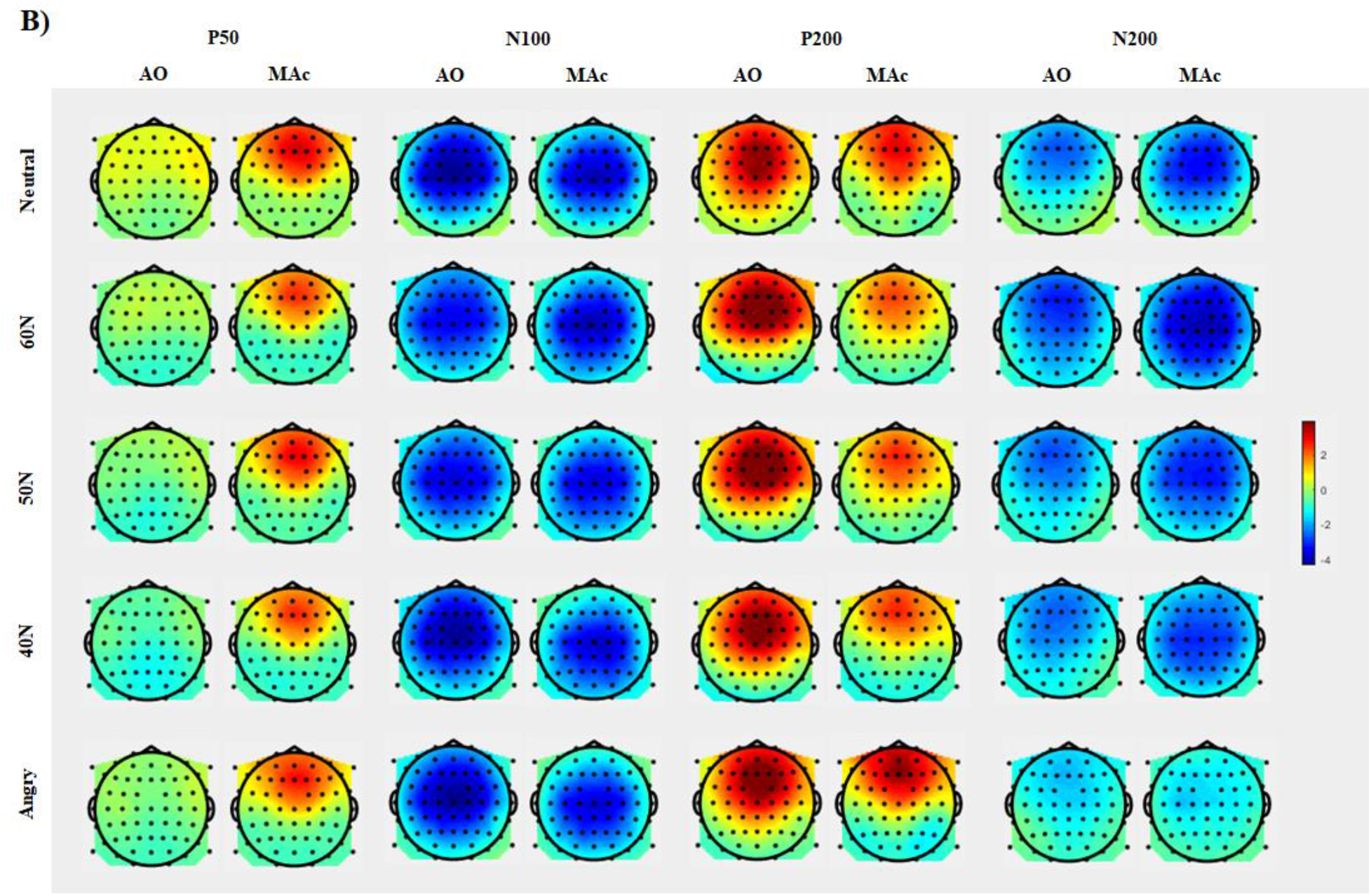
A) Grand average waveforms ± variance comparing self-generated and externally-generated for five types of self-voice originating from a fronto-central ROI. B) Topographic maps show voltage distribution in the ERP time window. Abbreviations: AO = Auditory Only; MAc = Motor Auditory corrected

### Statistical analyses

Statistical analyses of the ERP data were performed in R version 4.2.2 (2022-10-31) Copyright (C) 2022, using linear mixed models (LMMs) using the lmer and lmerTest packages [79, 80]. LMMs were used to control for random effects of participants influencing the outcome measure. Additionally, as HP measured by the LSHS is a continuous variable, LMMs were considered more appropriate than classical ANOVA to analyze the impact of HP on sensory feedback (condition) and emotional quality (stimulus type). Mean amplitude values of the ERPs were used as outcome measures, participants were used as random effects, and condition (2 levels: MAc and AO), stimulus type (5 levels: 100% neutral, 60-40% neutral-angry, 50-50% neutral-angry, 40-60% neutral-angry, 100% angry) and LSHS total or LSHS AVH (supplementary document A3) scores (continuous variable) were included as fixed effects, respectively, in the hypothesized models. For all models, the Gaussian distribution of model residuals and quantile-quantile plots confirmed their respective adequacy.

## 3. Results

Following our hypotheses, we probed the interaction of sensory feedback processing (conditions: MAc and AO), self-voice quality (stimulus type: 100% neutral, 60-40%: neutral-angry; 50-50%: neutral-angry; 40-60%: neutral-angry and 100% angry), and HP for each ERP. However, we did not find a significant interaction of sensory feedback processing and self-voice quality, therefore we reduced the models to probing the interaction of sensory feedback processing and self-voice quality with HP, respectively.

### N100 (primary outcome)

To probe the influence of HP based on LSHS total scores on condition and stimulus type, we tested [m1_N100 <-lmer(N100 ∼ + LSHS total * Condition + LSHS_total* Stimulus Type + (1|ID), data=data, REML = FALSE)] against the null model [m0_N100 <-lmer(N100 ∼ + (1|ID), data=data, REML = FALSE); AIC = 1356.5], which showed the best goodness of fit and yielded a significant difference (χ2(11) = 110.5, p = 0.000; AIC = 1268.0) (table 1, figure 3). We report a significant difference between self-generated (MAc) and externally-generated (AO) self-voices, regardless of the stimulus quality and HP. Moreover, the N100 suppression effect (AO minus MAc) decreased as HP increased.

**Figure 3:**
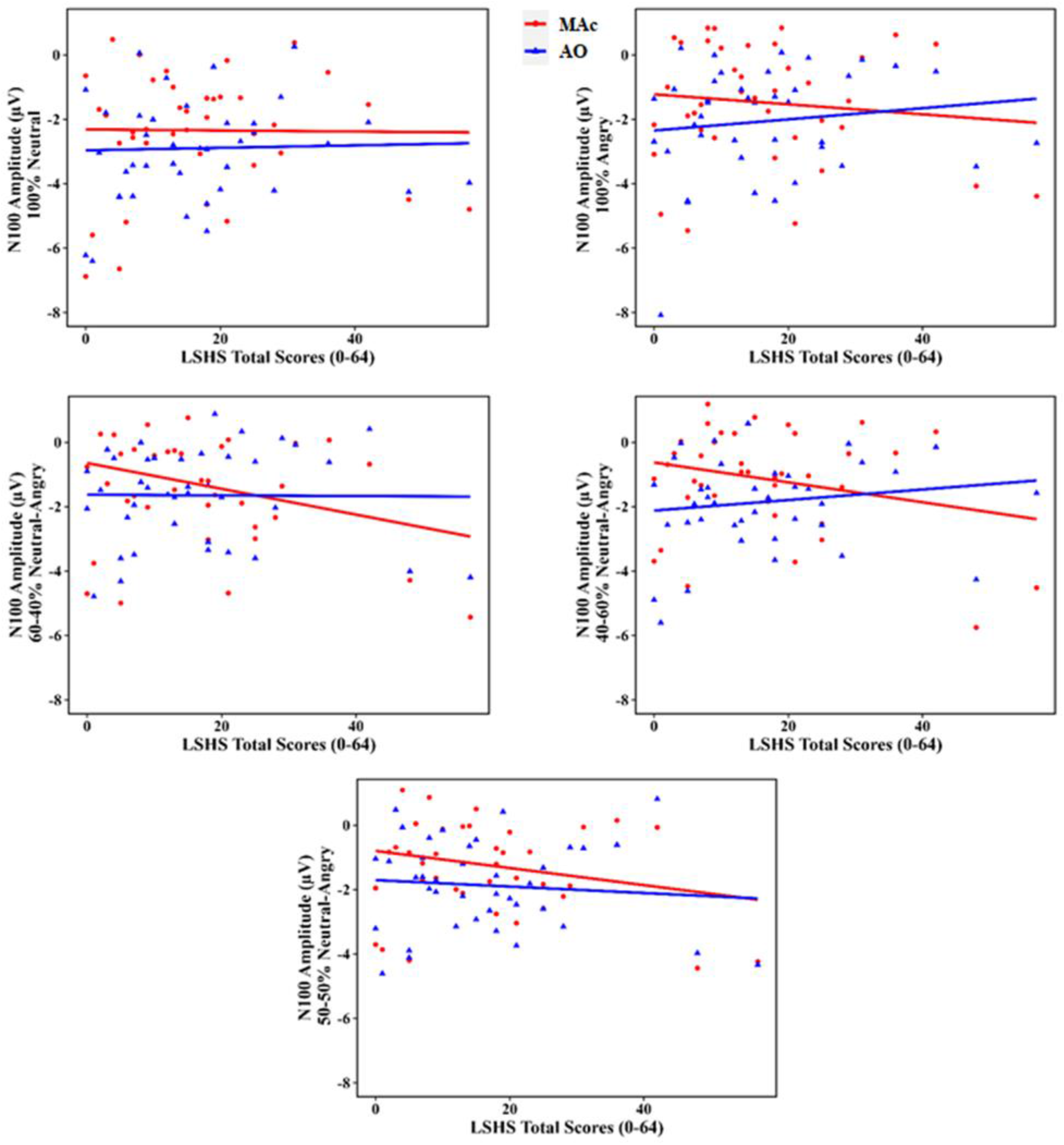
Scatter plots depicting the change in N100 amplitudes as a function of HP based on LSHS total scores for each stimulus type. The N100 suppression effect - difference in the N100 amplitude between AO and MAc, either decreased or reversed with increase in HP scores. Abbreviations: AO = Auditory Only; MAc = Motor Auditory corrected

**Table 1:** The linear mixed effects model of the N100 including the effect of HP based on LSHS total scores. Abbreviation: AO = Auditory Only; SE = standard error; SD = standard deviation; *p < 0.05; **p < 0.01; ***p < 0.001. Degrees of freedom for Fixed Effects: df = 360.00 (except Intercept: df = 64.18).

| Variable | Estimate | SE | t value | Pr(>\|t\|) |
| --- | --- | --- | --- | --- |
| **Fixed Effects** |  |  |  |  |
| Intercept | -2.121e+00 | 3.877e-01 | -5.471 | 7.88e-07 *** |
| AO | -1.031e+00 | 1.603e-01 | -6.433 | 3.98e-10 *** |
| LSHS total | -1.301e-02 | 1.847e-02 | -0.704 | 0.483998 |
| 60N | 1.502e+00 | 2.535e-01 | 5.926 | 7.27e-09 *** |
| 50N | 1.387e+00 | 2.535e-01 | 5.473 | 8.32e-08 *** |
| 40N | 1.265e+00 | 2.535e-01 | 4.992 | 9.32e-07 *** |
| Angry | 8.526e-01 | 2.535e-01 | 3.364 | 0.000852 *** |
| AO*LSHS total | 2.823e-02 | 7.640e-03 | 3.695 | 0.000254 *** |
| 60N*LSHS total | -2.161e-02 | 1.208e-02 | -1.789 | 0.074404 . |
| 50N*LSHS total | -1.938e-02 | 1.208e-02 | -1.605 | 0.109467 |
| 40N*LSHS total | -8.364e-03 | 1.208e-02 | -0.692 | 0.489133 |
| Angry*LSHS total | -2.002e-04 | 1.208e-02 | -0.017 | 0.986783 |
| **Groups** | **Name** | **Variance** | **SD** |  |
| **Random Effects** |  |  |  |  |
| Subjects | Intercept | 1.6894 | 1.2998 |  |
| Residual |  | 0.9714 | 0.9856 |  |
| Number of observations: 400, Subjects: 40 |  |  |  |  |

### P200 (secondary outcome)

To probe the influence of HP based on LSHS total scores on condition and stimulus type, we tested [m1_P200 <- lmer(P200 ∼ + LSHS total * Condition + LSHS_total* Stimulus Type + (1|ID), data=data, REML = FALSE)] against the null model [m0_P200 <- lmer(P200 ∼ + (1|ID), data=data, REML = FALSE); AIC = 1607.2], which showed the best goodness of fit and yielded a significant difference (χ2(11) = 147.82, p = 0.000; AIC = 1481.3) (table 2, figure 4). We report a significant difference between self-generated (MAc) and externally-generated (AO) self-voices, regardless of the stimulus quality and HP. Moreover, the P200 suppression effect (AO minus MAc) increased as HP levels increased.

**Figure 4:**
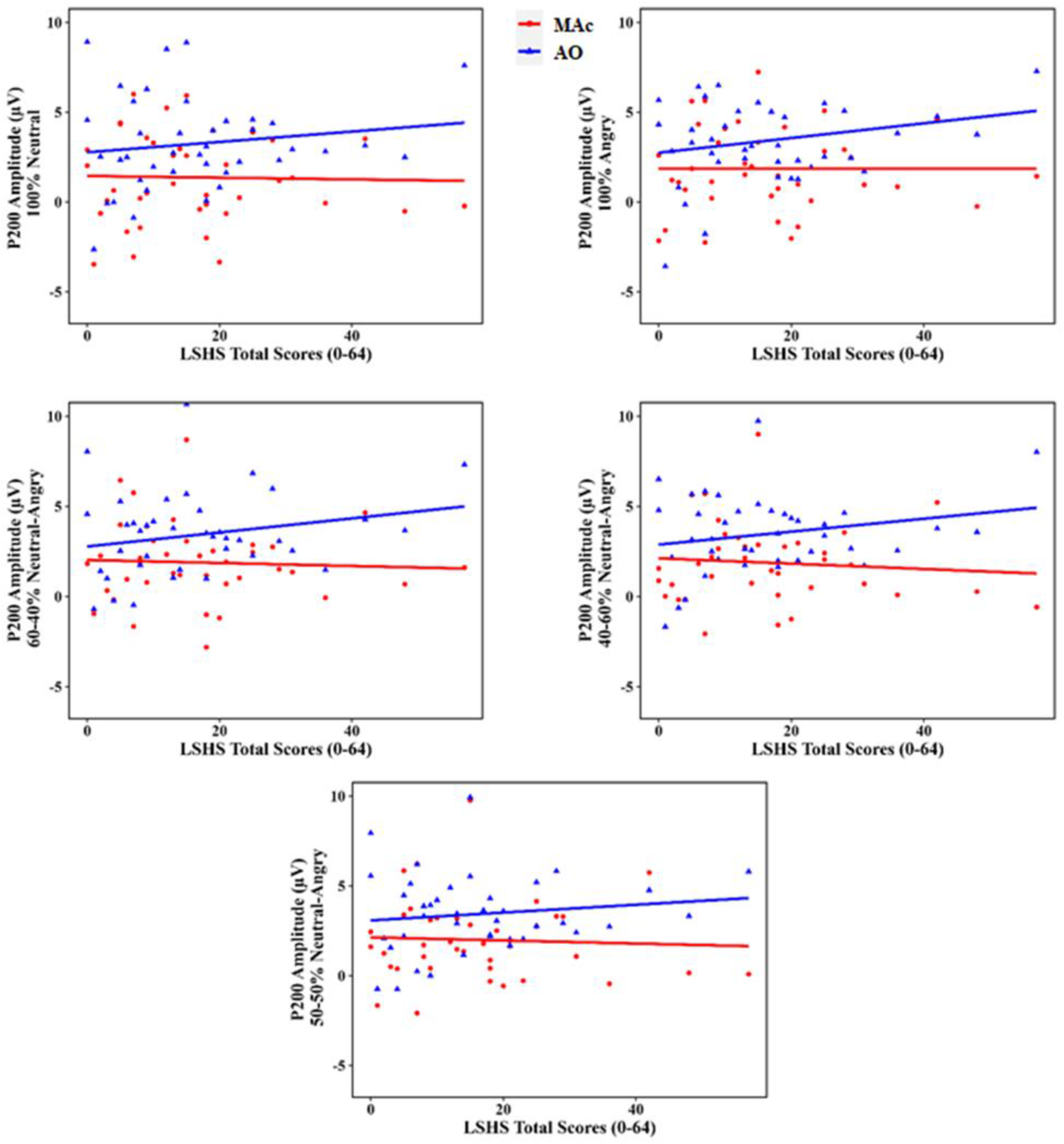
Scatter plots depicting the change in P200 amplitude as a function of HP based on LSHS total scores for each stimulus type. The P200 suppression effect - the difference in the amplitudes of AO and MAc, was modulated by HP such that the P200 amplitude for externally-generated (AO) voices increased with increase in HP scores. Abbreviations: AO = Auditory Only; MAc = Motor Auditory corrected

**Table 2:** The linear mixed effects model of the P200, including the effect of HP based on LSHS total scores. Abbreviation: AO = Auditory Only; SE = standard error; SD = standard deviation; *p < 0.05; **p < 0.01; ***p < 0.001. Degrees of freedom for Fixed Effects: df = 360.00 (except Intercept: df = 58.98).

| Variable | Estimate | SE | t value | Pr(>\|t\|) |
| --- | --- | --- | --- | --- |
| **Fixed Effects** |  |  |  |  |
| Intercept | 1.643231 | 0.547489 | 3.001 | 0.00394 ** |
| AO | 0.933481 | 0.206973 | 4.510 | 8.78e-06 *** |
| LSHS total | -0.008218 | 0.026090 | -0.315 | 0.75389 |
| 60N | 0.295346 | 0.327254 | 0.902 | 0.36740 |
| 50N | 0.497875 | 0.327254 | 1.521 | 0.12904 |
| 40N | 0.394437 | 0.327254 | 1.205 | 0.22888 |
| Angry | 0.187815 | 0.327254 | 0.574 | 0.56638 |
| AO*LSHS total | 0.040584 | 0.009863 | 4.115 | 4.81e-05 *** |
| 60N*LSHS total | 0.003236 | 0.015595 | 0.208 | 0.83574 |
| 50N*LSHS total | -0.005438 | 0.015595 | -0.349 | 0.72753 |
| 40N*LSHS total | -0.001537 | 0.015595 | -0.099 | 0.92153 |
| Angry*LSHS total | 0.008415 | 0.015595 | 0.540 | 0.58983 |
| **Groups** | **Name** | **Variance** | **SD** |  |
| **Random Effects** |  |  |  |  |
| Subjects | Intercept | 3.560 | 1.887 |  |
| Residual |  | 1.619 | 1.272 |  |
| Number of observations: 400, Subjects: 40 |  |  |  |  |

## Exploratory analysis

### P50

To probe the influence of HP (based on LSHS total scores) on condition and stimulus type, we tested [m1_P50 <- lmer(P50 ∼ + LSHS total * Condition + LSHS_total* Stimulus Type + (1|ID), data=data, REML = FALSE)] against the null model [m0_P50 <- lmer(P50 ∼ + (1|ID), data=data, REML = FALSE); AIC = 1423.0], which showed the best goodness of fit and yielded a significant difference (χ2(11) = 170.4, p = 0.000; AIC = 1274.6) (table 3, figure 5). Regardless of HP and stimulus type, there is a significant difference in the P50 response between self-generated (MAc) and externally-generated (AO) conditions. Further, the P50 suppression effect increased with increase in HP.

**Figure 5:**
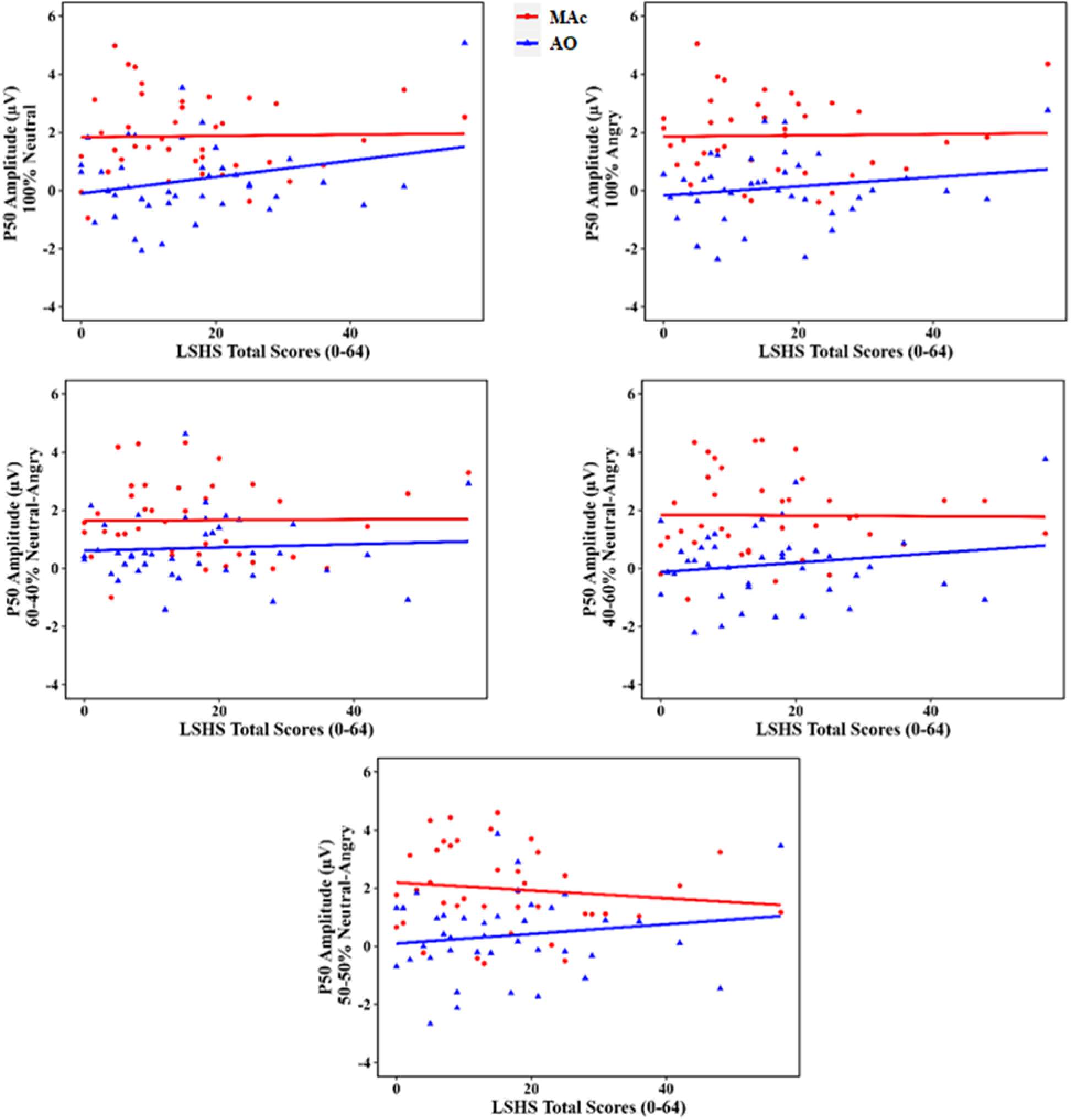
Scatter plots depicting the change in P50 as a function of HP (based on LSHS total scores) for each stimulus type. The P50 amplitude differences between AO and MAc decreased with increase in HP suggesting decrease in vigilance or sustained attention. Abbreviations: AO = Auditory Only; MAc = Motor Auditory corrected

**Table 3:** The linear mixed effects model of the P50 including the effect of HP based on LSHS total scores. Abbreviations: AO = Auditory Only; SE = standard error; SD = standard deviation; *p < 0.05; **p < 0.01; ***p < 0.001. Degrees of freedom for Fixed Effects: df = 360.00 (except Intercept: df = 116.24).

| Variable | Estimate | SE | t value | Pr(>\|t\|) |
| --- | --- | --- | --- | --- |
| **Fixed Effects** |  |  |  |  |
| Intercept | 1.774297 | 0.289666 | 6.125 | 1.27e-08 *** |
| AO | -1.810581 | 0.170145 | -10.641 | < 2e-16 *** |
| LSHS total | 0.006003 | 0.013804 | 0.435 | 0.664 |
| 60N | 0.256733 | 0.269023 | 0.954 | 0.341 |
| 50N | 0.275914 | 0.269023 | 1.026 | 0.306 |
| 40N | -0.012501 | 0.269023 | -0.046 | 0.963 |
| Angry | -0.025393 | 0.269023 | -0.094 | 0.925 |
| AO*LSHS total | 0.018250 | 0.008108 | 2.251 | 0.025 * |
| 60N*LSHS total | -0.011790 | 0.012820 | -0.920 | 0.358 |
| 50N*LSHS total | -0.013612 | 0.012820 | -1.062 | 0.289 |
| 40N*LSHS total | -0.007643 | 0.012820 | -0.596 | 0.551 |
| Angry*LSHS total | -0.006264 | 0.012820 | -0.489 | 0.625 |
| **Groups** | **Name** | **Variance** | **SD** |  |
| **Random Effects** |  |  |  |  |
| Subjects | Intercept | 0.6121 | 0.7823 |  |
| Residual |  | 1.0943 | 1.0461 |  |
| Number of observations: 400, Subjects: 40 |  |  |  |  |

### N200

To probe the influence of HP (based on LSHS total scores) on condition and stimulus type, we tested [m1_N200 <- lmer(N200 ∼ + LSHS total * Condition + LSHS_total* Stimulus Type + (1|ID), data=data, REML = FALSE)] against the null model [m0_N200 <- lmer(P200 ∼ + (1|ID), data=data, REML = FALSE); AIC = 1388.4], which showed the best goodness of fit and yielded a significant difference (χ2(11) = 106.66, p = 0.000; AIC = 1303.7) (table 4, figure 6). There was no significant interaction of HP with condition (MAc or AO) or stimulus type (5 types of self-voice) and no main effect of condition (MAc/AO). However, there was a main effect of stimulus type for 50-50% neutral-angry, 40-60% neutral-angry and 100% angry self-voice.

**Figure 6:**
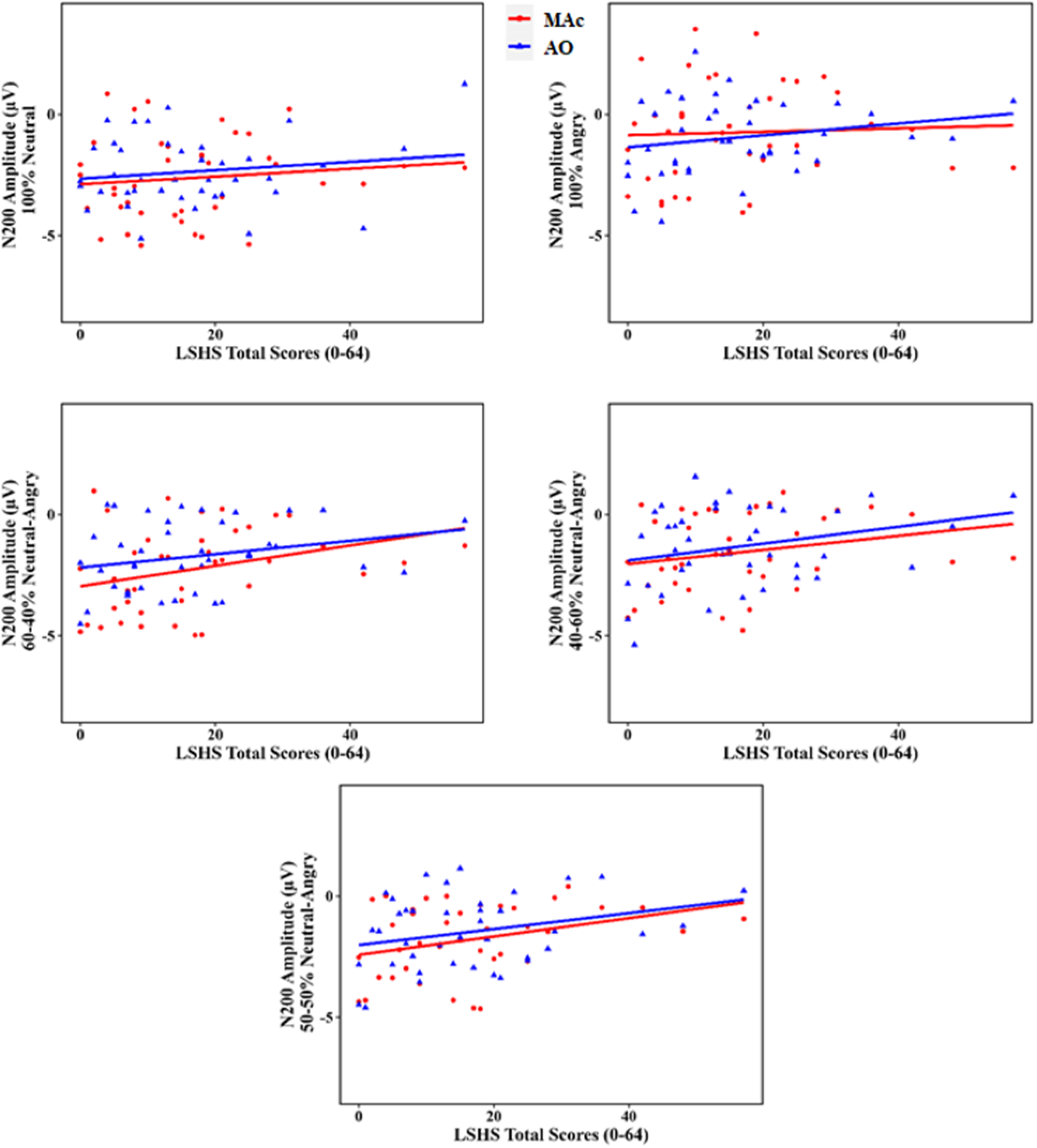
Scatter plots depicting the change in N200 amplitudes as a function of HP (based on LSHS total scores) for each stimulus type. No difference between the N200 amplitudes for AO and MAc. Abbreviations: AO = Auditory Only; MAc = Motor Auditory corrected

**Table 4:** The linear mixed effects model of the N200, including the effect of HP based on LSHS total scores. Abbreviations: AO = Auditory Only; SE = standard error; SD = standard deviation; *p < 0.05; **p < 0.01; ***p < 0.001. Degrees of freedom for Fixed Effects: df = 360.00 (except Intercept: df =75.49).

| Variable | Estimate | SE | t value | Pr(>\|t\|) |
| --- | --- | --- | --- | --- |
| **Fixed Effects** |  |  |  |  |
| Intercept | -2.874e+00 | 3.617e-01 | -7.946 | 1.45e-11 *** |
| AO | 2.116e-01 | 1.706e-01 | 1.241 | 0.2156 |
| LSHS total | 1.610e-02 | 1.724e-02 | 0.934 | 0.3531 |
| 60N | 1.943e-01 | 2.697e-01 | 0.721 | 0.4716 |
| 50N | 5.470e-01 | 2.697e-01 | 2.028 | 0.0433 * |
| 40N | 7.978e-01 | 2.697e-01 | 2.958 | 0.0033 ** |
| Angry | 1.663e+00 | 2.697e-01 | 6.165 | 1.89e-09 *** |
| AO*LSHS total | 9.931e-04 | 8.128e-03 | 0.122 | 0.9028 |
| 60N*LSHS total | 1.821e-02 | 1.285e-02 | 1.417 | 0.1573 |
| 50N*LSHS total | 1.881e-02 | 1.285e-02 | 1.463 | 0.1443 |
| 40N*LSHS total | 1.533e-02 | 1.285e-02 | 1.193 | 0.2337 |
| Angry*LSHS total | -9.401e-04 | 1.285e-02 | -0.073 | 0.9417 |
| **Groups** | **Name** | **Variance** | **SD** |  |
| **Random Effects** |  |  |  |  |
| Subjects | Intercept | 1.318 | 1.148 |  |
| Residual |  | 1.100 | 1.049 |  |
| Number of observations: 400, Subjects: 40 |  |  |  |  |

### Correlational analysis

If the N100 response is sensitive to discrepancies in sensory feedback processing and likely engages increased attentional resource allocation in self-voice production, we would expect corresponding changes at subsequent perceptual processing stages, like the N200, associated with error awareness and attentional control [76, 81–83]. Therefore, to examine a potential relationship between N100 and N200, we correlated their amplitudes for each condition (AO and MAc) per stimulus type. Significant correlation between N100 and N200 mean amplitudes were observed for certain but not for uncertain (60-40% neutral-angry and 50-50% neutral-angry) self-voice processing.

## 4. Discussion (1427)

The fundamental mechanisms that contribute to perceiving voices without any external sensory input remain inadequately understood. By varying the self-voice emotional quality, this study investigated whether changes in certainty of sensory feedback during self-voice production are linked to an individual’s proneness to experience hallucinations and attentional allocation. The ERPs evoked by an individual’s self- and externally-generated self-voice were examined within a classical auditory-motor paradigm (figure 1; [39]). As expected, the N100 and P200 were suppressed for self-compared to externally-generated self-voices (table 1, 2; figure 2, 3, 4). Disengagement of resource allocation was observed for self-voice stimuli with ambiguous emotional quality, but not for those with unambiguous emotional quality (figure 7), suggesting that ambiguous self-voice stimuli did not recruit additional resources for further processing. HP modulated the P50, N100 and P200 suppression effects (table 1, 2, 3; figure 3, 4, 5), indicating increased error awareness and attention allocation in high HP individuals in self-voice processing likely stemming from alterations in sensory feedback processing, and/or attentional control.

**Figure 7:**
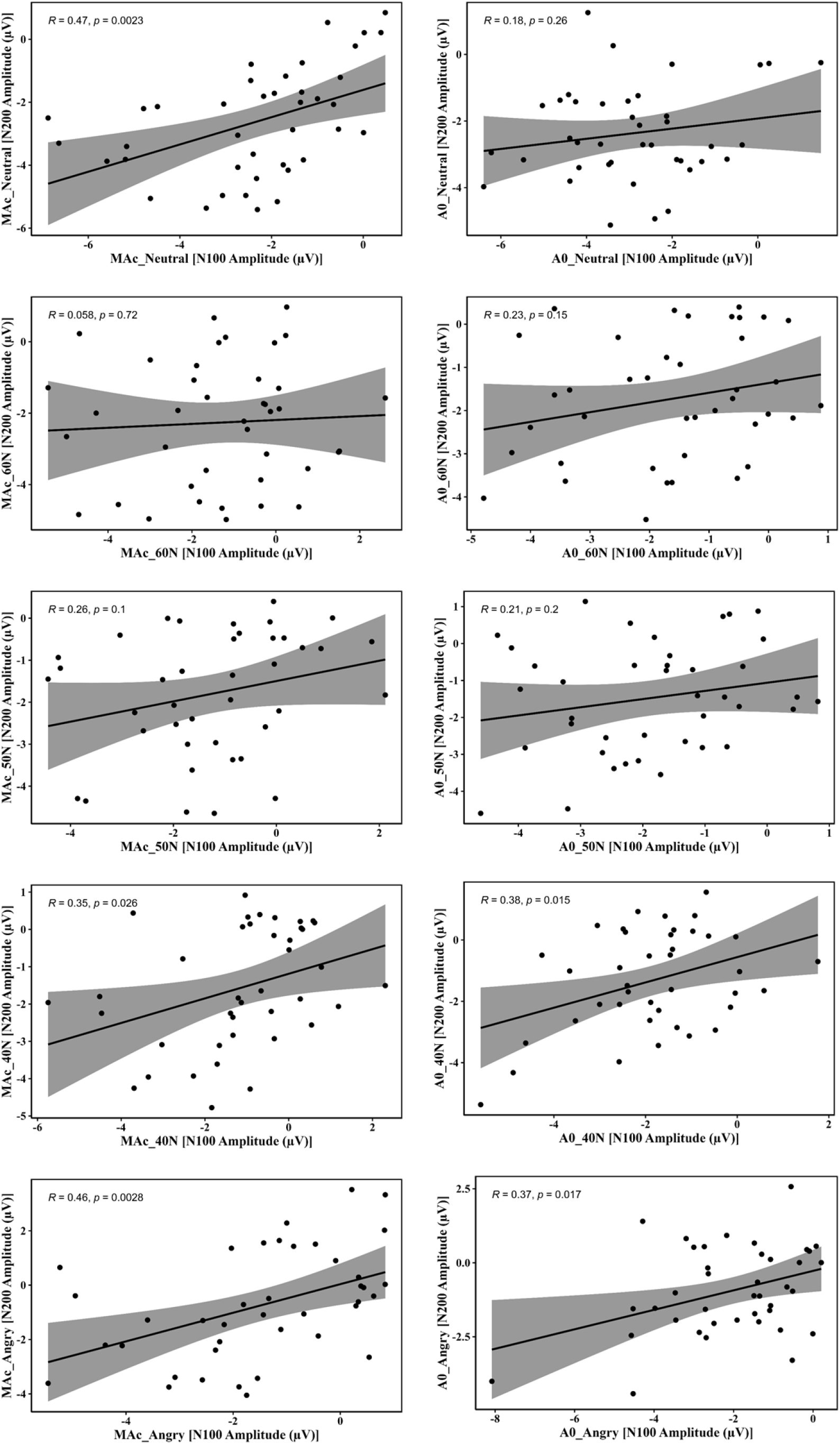
Correlations between N100 and N200 mean amplitudes for MAc and AO per stimulus type. The significant correlations between N100 and N200 amplitudes for certain but not uncertain self-voice may suggest disengagement of, or reduced resource allocation for uncertain self-voices (60-40% neutral-angry and 50-50% neutral-angry), possibly due to perceiving them as dissimilar to one’s own voice. Abbreviations: AO = Auditory Only; MAc = Motor Auditory corrected

### Sensory feedback processing and attentional control

Principally, the P50, N100 as well as P200 and N200 responses can be indicative of expected sound input albeit at different information processing stages (P50 and N100: sensory feedback processing, attention allocation [33, 34, 84, 85], P200: conscious distinction between self- and externally-generated voice [72–74, 86], N200: error awareness/monitoring and attentional control; [86–89]). We report a more enhanced P50 response to self-than externally-generated voices, regardless of the self-voice quality or HP (table 3, figure 5). This is in contrast to previous findings that reported P50 suppression for self-generated tones [84]. However, the P50 suppression was also associated with the decline in attention and vigilance over the course of the task [70, 90]. The enhanced P50 response to self-generated voices may likely reflect increased vigilance and sustained attention [90] towards the unexpected voice qualities of the self-generated voices. The self-generated condition preceded the externally-generated condition in the current task-design, which means that the novelty of unexpectedness of the self-voice remained higher in the self-generated than the passive listening condition (externally-generated voice). This interpretation is supported by studies reporting an association between the enhanced P50 and deviance detection and the encoding of irregularities in the auditory environment [91]. Of note, we report the opposite for the N100 response, i.e., reduced amplitudes for self-compared to the externally-generated voice, independent of the voice quality (neutral-angry) (figure 2; [39, 74]). This suppression likely relates to the reduction of neural activity in the AC suggesting a match between predicted and perceived sensory feedback of the self-voice. The explanation for the reversed P50 and N100 effects for the self-generated voice likely lies in the early time window (40-60 ms), which remains unaffected by expectation, while the later time window (100-200 ms) is impacted [92]. Further, we report a significant correlation between the N100 and N200 amplitudes for certain but not uncertain/ambiguous (60-40% neutral-angry, 50-50% neutral-angry) self-voices (figure 7). This may indicate reduced resource allocation and limited processing due to disengagement of attentional resources for non-self-relevant voices as these uncertain/ambiguous voices were not entirely perceived as either purely self-voice or entirely someone else’s voice. Nonetheless, the lack of an interaction between condition and self-voice quality (based on N100 or N200 responses; table 1, table 4) and “own-ness” rating (supplementary figure 2) indicates the possibility that the differences between the uncertain/ambiguous and certain/unambiguous voices were still within the acceptable range of feasible physiological change in voice quality and thus might not have caused big enough sensory perturbations or mismatch between predicted and perceived feedback.

### Hallucination proneness, sensory feedback processing, and attention allocation

HP modulated P50, N100, and P200 effects. As HP (based on LSHS total scores) increased from low to high, the P50 response for self-compared to externally-generated voices decreased (table 3, figure 5). This reduction of the P50 response might reflect decreased vigilance and sustained attention due to complexity of sensory information, that is, the variation in emotional quality as well as recognition of self-voice [90]. Individuals who score high in HP have been reported to show impaired filtering of information, thus might not be able to separate relevant from irrelevant stimuli, which leads to sensory overload [93–96] Similarly, the N100 suppression effect was reduced such that the N100 response for the self-generated voice is increased with increased HP (based on both LSHS total [table 1, figure 3] and AVH scores [see supplementary document]). This change in N100 suppression likely indicates that there is a mismatch between the predicted and perceived sensory feedback in high HP individuals rooted in altered sensory feedback processing [39, 97]. The N100 response is also associated with spontaneous attention allocation [33, 34, 98]. Attention and prediction show complementary effects on the N100 response: the N100 amplitude decreases with an increase in predictability whereas it increases when more attention needs to be allocated to an event [33, 34]. Therefore, an increased N100 amplitude in response to the self-generated voice with increased HP might also indicate altered error awareness and/or inability to suppress attention allocation to an irrelevant stimulus - one’s own self-generated voice. In sum, individuals with high HP display altered sensory feedback processing for the self-voice and might misattribute attentional resources to it. Both accounts are associated with theories such as self-monitoring and salience misattribution explaining AVH [9, 42, 43, 99, 100].

The P200 response is associated with a more conscious detection of sensory feedback to a self-generated compared to an externally-generated stimulus [72, 73, 86, 101, 102]. The current findings also confirm an increase in the P200 suppression effect with increased P200 response for externally-generated voices with an increase in HP (based on both LSHS total and AVH scores) (table 2; figure 4). The P200 response is reportedly more sensitive to temporal predictability of an auditory stimulus [39, 78, 103–105]. Specifically, increased P200 responses were reported with longer delays between a button-press and sensory feedback, reflecting a potential decrease in the sense of agency when expectations are not matched immediately [39, 78, 103, 105]. As the timing of the different self-voices were unpredictable in the externally-compared to the self-generated condition, an enhanced P200 response in high HP during passive listening to self-voice could merely reflect more mindful processing of the five types of voices with variable onsets, triggering more attentional resources than in low HP.

The current sample of participants consisted of non-voice hearers from the general population who varied in HP (N = 38) as well as 2 voice hearers with a psychotic disorder. Given the sample size calculations (supplementary document A2) and early termination of the study, the current had limited statistical power, which may additionally explain the absence of an observed three-way interaction between conditions (AO, MAc), voice quality (5 types of self-voice), and HP. Furthermore, because of the small number of voice hearers with a psychotic disorder we were unable fully to explore the putative psychosis continuum. Previous studies probing prediction of sensory feedback to the self-voice reported no N100 suppression effect in voice hearers with a psychotic disorder [106, 107] whereas non-clinical voice hearers or high HP individuals showed a reversed N100 suppression effect [38, 39, 108]. These studies either used group differences [107] or hypothesized a linear relationship between HP and sensory feedback processing/attentional control, measured by the N100 suppression effect, using LMMs [39, 108]. Future studies should consider that individuals on the postulated HP severity continuum (voice hearers with psychotic disorder, at high-risk of psychosis/prodromal and non-clinical voice hearers) may show a nonlinear relationship with the N100 suppression effect. Future studies should also analyze the influence of HP on pre-button-press neural activity, including the readiness potential [109, 110] and pre-stimulus alpha power, which have both been related to the prediction of the sensory consequences of a self-generated stimulus [38, 39].

Using an established experimental paradigm in combination with a manipulation of the emotional quality of the self-voice, we detected associations between HP and electrophysiological indicators of sensory feedback during self-voice production. The findings contribute to a better understanding of the association between predicted and perceived sensory feedback and its potential role in auditory hallucinations.

## Funding

The current project was supported by internal funds.

## Data availability

The data and analysis pipelines that support the findings of this study are available from the corresponding author upon reasonable request.

## Author Contributions

SXD, MS, AP, DL, TvA, SK conceptualized the project. SXD, AP, MS, SK designed the experiment, SXD prepared materials, SXD and HH collected data, SXD analyzed the data, and wrote the first draft of the manuscript and SXD, HH, MS, TvA, DL, AP, SK edited and refined the manuscript. All authors have approved the final version of the manuscript.

## Competing Interests

The authors declare that they have no competing interests.

## Acknowledgements

The authors would like to thank our interns Alex Kalberer and Maren Cremer for their support with data collection.

## Notes

### Competing Interest Statement

The authors have declared no competing interest.

