## supplementary document for "Exploring Neural Dynamics in Self-Voice Processing and Perception: Implications for Hallucination Proneness"

**Section A**

**A1. Exclusion criteria:** Participants were excluded if they (i) had undergone any previous neurosurgery or had a neurological disorder, (ii) refused to participate in the EEG, (iii) were unable to fully comprehend the purpose of the study or to make a rational decision whether or not to participate, (iv) heard voices exclusively due to substance abuse (drug or alcohol addiction).

**A2. Sample size calculation:**

As the primary objective was to examine N100 suppression effects (AO - MAc) as a function of hallucination proneness (HP), the total number of required participants was based on a power analysis for the correlation between the quantitative covariate (HP) and a linear contrast of the within subjects measure (e.g., N100 suppression effect for self-voice changing from fully neutral to angry in five steps) using G*Power statistical software (3.1.9) [1] with a detectable effect size of (d) = 0.50 resulting in ρ of 0.24, α = 0.05, power (1-β) = 0.80 and adjusting for confounding factors (e.g., age, education, gender, medication). Thus, the final sample size was 210 participants in total including voice hearers and participants from the general population. Based on previous literature, 7-13% of the general population experience auditory verbal hallucinations (AVH) [2-4]. Thus, 16-30 voice hearers (belonging to the three categories of non-voice hearers, non-clinical voice hearers and voice hearers with psychotic disorder) were recruited. Considering the lower prevalence of voice hearers in the general population, participants were recruited from the general population through advertisements in newspaper and social media, as well as voice hearing centers.

*t tests - Correlation: Point biserial model*

Analysis: A priori: Compute required sample size

Input: Tail(s) = Two

Effect size |ρ| = 0.24

α err prob = 0.05

Power (1-β err prob) = 0.80

Output: Noncentrality parameter δ = 2.8296274

Critical t = 1.9785245

Df = 129

Total sample size = 131

Actual power = 0.8019318

After adjusting for the potentially confounding factors such as age, gender and education, the sample size is multiplied by 1.5 (variance inflation factor; VIF), 1.05 (to account for the power loss due to interim analysis) and approximately 10% drop out rate = (131*1.5*1.07) + 21 = 231.

The current study was concluded prematurely due to slow recruitment of voice hearers, leading to a final sample of 45 recruited participants.

**A3. Results**

*HP based on LSHS AVH scores*

P50: To probe the influence of HP ( based on LSHS AVH scores) on condition and stimulus type, we tested [m1_P50 <- lmer(P50 ~ + LSHS AVH * Condition + LSHS AVH * Stimulus Type + (1|ID), data=data, REML = FALSE) ] against the null model [m0_P50 <- lmer(P50 ~ + (1|ID), data=data, REML = FALSE); AIC = 1423.0], which showed the best goodness of fit and yielded a significant difference (χ2(11) = 167.77, p = 0.000; AIC = 1277.3) (supplementary table 2 and figure 5).

N100: To probe the influence of HP (based on LSHS AVH scores) on condition and stimulus type, we tested [m1_N100 <- lmer(N100 ~ + LSHS AVH * Condition + LSHS AVH * Stimulus Type + (1|ID), data=data, REML = FALSE) ] against the null model [m0_N100 <- lmer(N100 ~ + (1|ID), data=data, REML = FALSE); AIC = 1356.5], which showed the best goodness of fit and yielded a significant difference (χ2(11) = 110.5, p = 0.000; AIC = 1268.0) (supplementary table 3 and figure 6).

P200: To probe the influence of HP (based on LSHS AVH scores) on condition and stimulus type, we tested [m1_P200 <- lmer(P200 ~ + LSHS AVH * Condition + LSHS AVH * Stimulus Type + (1|ID), data=data, REML = FALSE) ] against the null model [m0_P200 <- lmer(P200 ~ + (1|ID), data=data, REML = FALSE); AIC = 1607.2], which showed the best goodness of fit and yielded a significant difference (χ2(11) = 150.59, p = 0.000; AIC = 1478.6) (supplementary table 4 and figure 7).

N200: To probe the influence of HP (based on LSHS AVH scores) on condition and stimulus type, we tested [m1_N200 <- lmer(N200 ~ + LSHS AVH * Condition + LSHS_AVH * Stimulus Type + (1|ID), data=data, REML = FALSE) ] against the null model [m0_N200 <- lmer(N200 ~ + (1|ID), data=data, REML = FALSE); AIC = 1388.4], which showed the best goodness of fit and yielded a significant difference (χ2(11) = 107.95, p = 0.000; AIC = 1302.5) (supplementary table 5, and figure 8).

**Section B**

**Tables and table legends**

**Supplementary table 1:** Neutral-angry continua with 11 voice morphs.

a)	Neutral-to-angry

| Emotion/Morphs | **1*** | 2 | 3 | 4 | **5*** | **6*** | **7*** | 8 | 9 | 10 | **11*** |
| --- | --- | --- | --- | --- | --- | --- | --- | --- | --- | --- | --- |
| Neutral | **100%** | 90% | 80% | 70% | **60%** | **50%** | **40%** | 30% | 20% | 10% | **0%** |
| Angry | **0%** | 10% | 20% | 30% | **40%** | **50%** | **60%** | 70% | 80% | 90% | **100%** |

*****final auditory stimuli

**Supplementary table 2:** Linear mixed effects model of P50 amplitude including the effect of HP based on LSHS AVH scores. Abbreviations: SE = standard error; SD = standard deviation; AO = auditory only; *p < 0.05; **p < 0.01; ***p < 0.001. Degrees of freedom for Fixed Effects: df = 360.00 (except Intercept: df = 118.89).

| **Variable** | **Estimate** | **SE** | **t value** | **Pr(>\|t\|)** |
| --- | --- | --- | --- | --- |
| **Fixed Effects** | | | | |
| Intercept | 1.786494 | 0.227496 | 7.853 | 2.01e-12 *** |
| AO | -1.637872 | 0.134772 | -12.153 | < 2e-16 *** |
| LSHS AVH | 0.034515 | 0.056391 | 0.612 | 0.542 |
| 60N | 0.137511 | 0.213094 | 0.645 | 0.519 |
| 50N | 0.108717 | 0.213094 | 0.510 | 0.610 |
| 40N | -0.124078 | 0.213094 | -0.582 | 0.561 |
| Angry | -0.133966 | 0.213094 | -0.629 | 0.530 |
| AO*LSHS total | 0.051218 | 0.033407 | 1.533 | 0.126 |
| 60N*LSHS total | -0.030061 | 0.052821 | -0.569 | 0.570 |
| 50N*LSHS total | -0.023005 | 0.052821 | -0.436 | 0.663 |
| 40N*LSHS total | -0.005906 | 0.052821 | -0.112 | 0.911 |
| Angry*LSHS total | 0.001943 | 0.052821 | 0.037 | 0.971 |
| **Groups** | **Name** | **Variance** | **SD** |  |
| **Random Effects** | | | | |
| Subjects | Intercept | 0.5963 | 0.7722 |  |
| Residual |  | 1.1048 | 1.0511 |  |
| Number of observations: 400, Subjects: 40 | | | | |

**Supplementary table 3:** Linear mixed effects model of N100 amplitude including the effect of HP based on LSHS AVH scores. Abbreviations: SE = standard error; SD = standard deviation, AO = auditory only; *p < 0.05; **p < 0.01; ***p < 0.001. Degrees of freedom for Fixed Effects: df = 360.00 (except Intercept: df = 65.29).

| **Variable** | **Estimate** | **SE** | **t value** | **Pr(>\|t\|)** |
| --- | --- | --- | --- | --- |
| **Fixed Effects** | | | | |
| Intercept | -2.000372 | 0.302946 | -6.603 | 8.52e-09 *** |
| AO | -0.842473 | 0.127317 | -6.617 | 1.33e-10 *** |
| LSHS AVH | -0.133046 | 0.075094 | -1.772 | 0.081107 . |
| 60N | 1.215397 | 0.201306 | 6.038 | 3.90e-09 *** |
| 50N | 1.083033 | 0.201306 | 5.380 | 1.34e-07 *** |
| 40N | 1.122067 | 0.201306 | 5.574 | 4.89e-08 *** |
| Angry | 0.761286 | 0.201306 | 3.782 | 0.000182 *** |
| AO*LSHS AVH | 0.110205 | 0.031559 | 3.492 | 0.000539 *** |
| 60N*LSHS AVH | -0.028115 | 0.049900 | -0.563 | 0.573486 |
| 50N*LSHS AVH | -0.006568 | 0.049900 | -0.132 | 0.895359 |
| 40N*LSHS AVH | 0.001936 | 0.049900 | 0.039 | 0.969067 |
| Angry*LSHS AVH | 0.034853 | 0.049900 | 0.698 | 0.485341 |
| **Groups** | **Name** | **Variance** | **SD** |  |
| **Random Effects** | | | | |
| Subjects | Intercept | 1.641 | 1.281 |  |
| Residual |  | 0.986 | 0.993 |  |
| Number of observations: 400, Subjects: 40 | | | | |

**Supplementary table 4:** Linear mixed effects model of P200 amplitude including the effect of HP based on LSHS AVH scores. Abbreviations: SE = standard error; SD = standard deviation, AO = auditory only; *p < 0.05; **p < 0.01; ***p < 0.001. Degrees of freedom for Fixed Effects: df = 360.00 (except Intercept: df = 58.65).

| **Variable** | **Estimate** | **SE** | **t value** | **Pr(>\|t\|)** |
| --- | --- | --- | --- | --- |
| **Fixed Effects** | | | | |
| Intercept | 1.81249 | 0.43259 | 4.190 | 9.52e-05 *** |
| AO | 1.16573 | 0.16246 | 7.175 | 4.15e-12 *** |
| LSHS AVH | -0.12090 | 0.10723 | -1.127 | 0.264 |
| 60N | 0.26002 | 0.25688 | 1.012 | 0.312 |
| 50N | 0.32058 | 0.25688 | 1.248 | 0.213 |
| 40N | 0.30127 | 0.25688 | 1.173 | 0.242 |
| Angry | 0.08711 | 0.25688 | 0.339 | 0.735 |
| AO*LSHS AVH | 0.17403 | 0.04027 | 4.321 | 2.01e-05 *** |
| 60N*LSHS AVH | 0.03520 | 0.06367 | 0.553 | 0.581 |
| 50N*LSHS AVH | 0.03457 | 0.06367 | 0.543 | 0.587 |
| 40N*LSHS AVH | 0.02682 | 0.06367 | 0.421 | 0.674 |
| Angry*LSHS AVH | 0.09504 | 0.06367 | 1.493 | 0.136 |
| **Groups** | **Name** | **Variance** | **SD** |  |
| **Random Effects** | | | | |
| Subjects | Intercept | 3.590 | 1.895 |  |
| Residual |  | 1.605 | 1.267 |  |
| Number of observations: 400, Subjects: 40 | | | | |

**Supplementary table 5:** Linear mixed effects model of N200 amplitude including the effect of HP based on LSHS AVH scores. Abbreviations: SE = standard error; SD = standard deviation, AO = auditory only; *p < 0.05; **p < 0.01; ***p < 0.001. Degrees of freedom for Fixed Effects: df = 360.00 (except Intercept: df = 72.86).

| **Variable** | **Estimate** | **SE** | **t value** | **Pr(>\|t\|)** |
| --- | --- | --- | --- | --- |
| **Fixed Effects** | | | | |
| Intercept | -2.54473 | 0.29093 | -8.747 | 5.64e-13 *** |
| AO | 0.08674 | 0.13378 | 0.648 | 0.51718 |
| LSHS AVH | -0.02490 | 0.07211 | -0.345 | 0.73091 |
| 60N | 0.24894 | 0.21153 | 1.177 | 0.24004 |
| 50N | 0.63322 | 0.21153 | 2.994 | 0.00295 ** |
| 40N | 0.87151 | 0.21153 | 4.120 | 4.70e-05 *** |
| Angry | 1.60508 | 0.21153 | 7.588 | 2.81e-13 *** |
| AO*LSHS AVH | 0.05596 | 0.03316 | 1.688 | 0.09237 . |
| 60N*LSHS AVH | 0.09777 | 0.05243 | 1.865 | 0.06306 . |
| 50N*LSHS AVH | 0.08913 | 0.05243 | 1.700 | 0.09003 . |
| 40N*LSHS AVH | 0.07129 | 0.05243 | 1.360 | 0.17481 |
| Angry*LSHS AVH | 0.01670 | 0.05243 | 0.318 | 0.75031 |
| **Groups** | **Name** | **Variance** | **SD** |  |
| **Random Effects** | | | | |
| Subjects | Intercept | 1.406 | 1.186 |  |
| Residual |  | 1.089 | 1.043 |  |
| Number of observations: 400, Subjects: 40 | | | | |

**Section C**

**Figures and Figure legends**

**Supplementary figure 1:** Schematic representing the paradigm design.

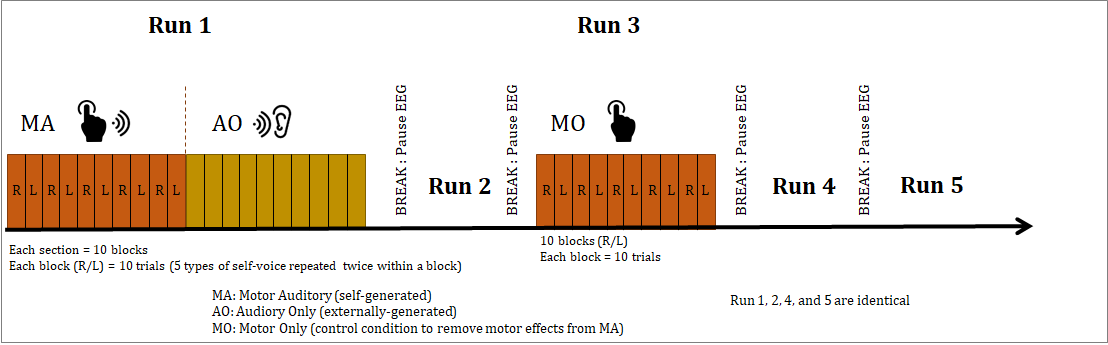

**Supplementary figure 2:** Post experiment stimuli rating. A) Arousal rating on a scale of 0-9 for each voice stimulus. B) Valence rating on a scale of 0-9 for each voice stimulus. C) Ownness rating on a scale of 0-10 for each voice stimulus.

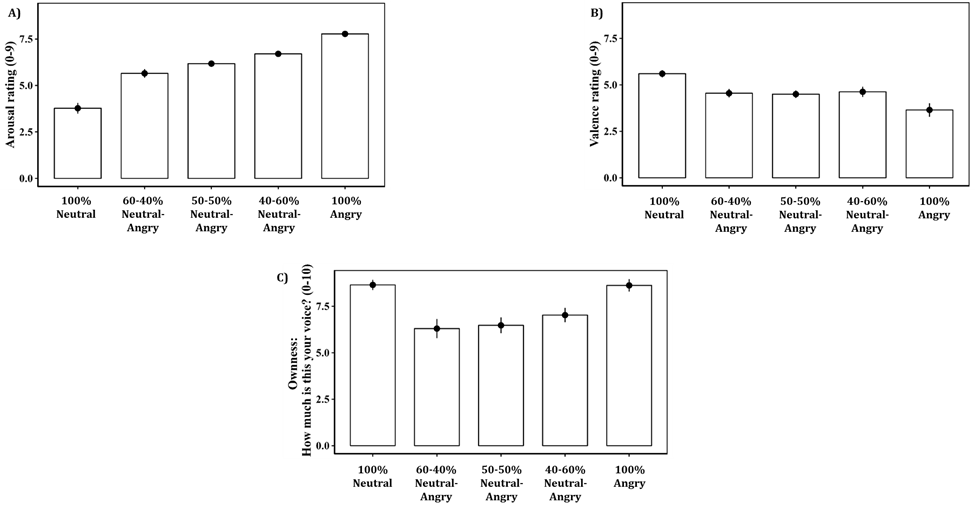

**Supplementary figure 3:** Mean ERP amplitudes for MAc and AO and suppression effects (AO - MAc) per stimulus. Abbreviations: AO = Auditory Only; MAc = Motor Auditory corrected.

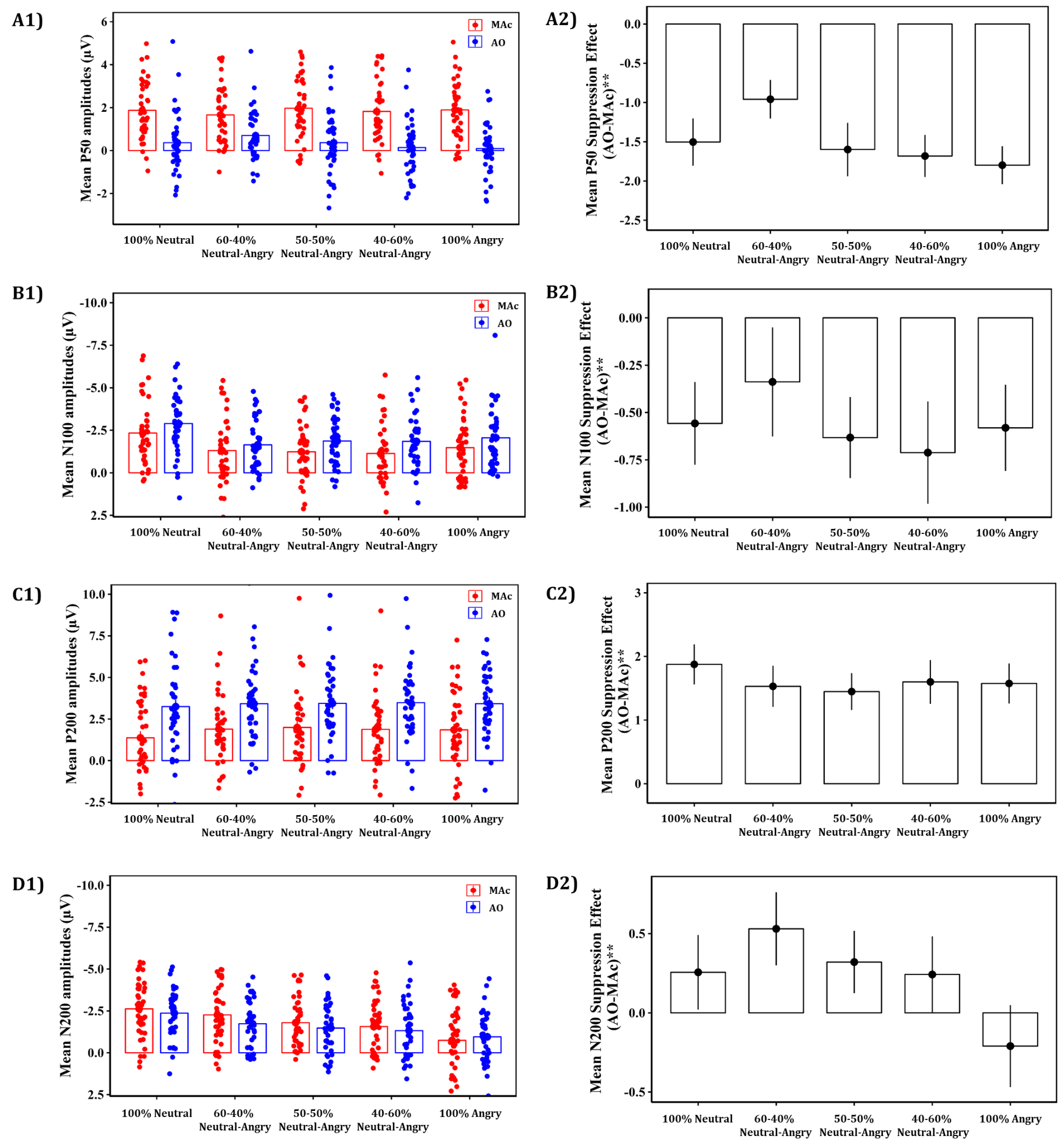

**Supplementary figure 4:** Scatter plots depicting the change in P50 amplitudes as a function of HP based on LSHS AVH scores for each stimulus type. Abbreviations: AO = Auditory Only; MAc = Motor Auditory corrected.

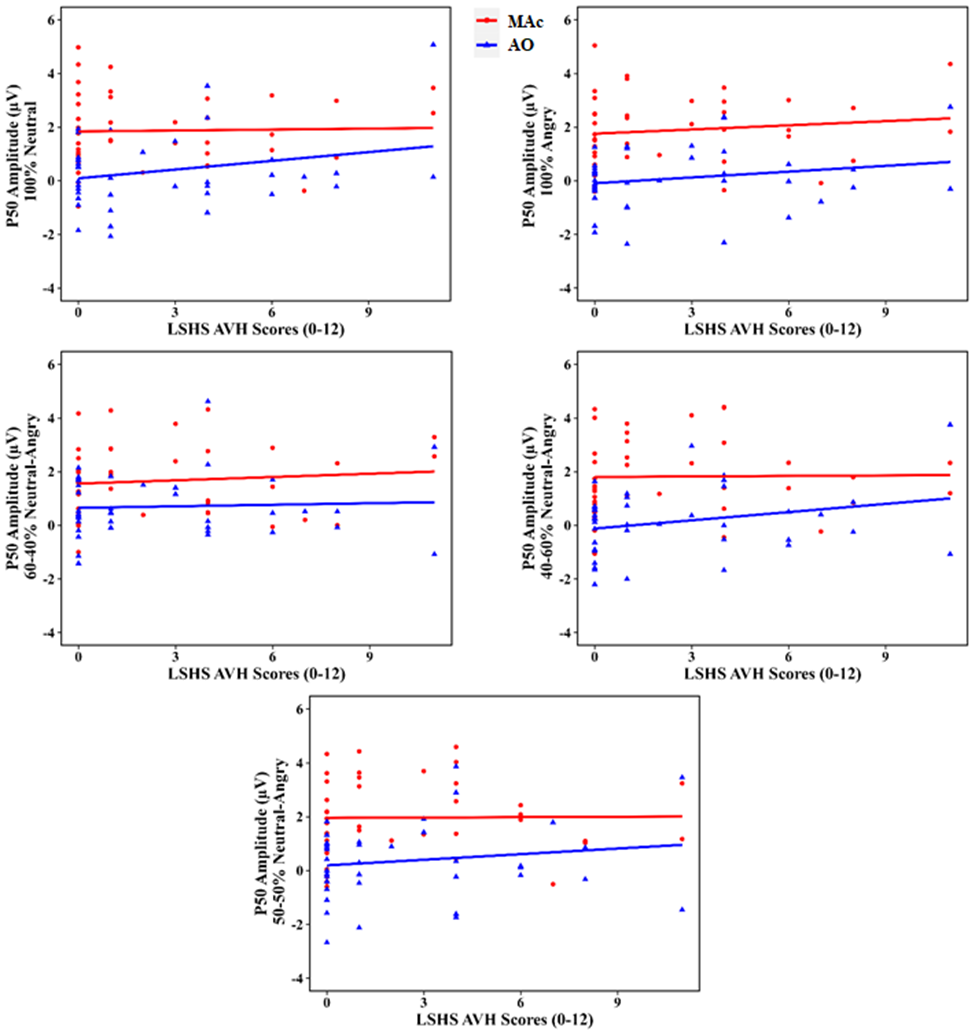

**Supplementary figure 5:** Scatter plots depicting the change in N100 amplitudes as a function of HP based on LSHS AVH scores for each stimulus type. Abbreviations: AO = Auditory Only; MAc = Motor Auditory corrected**.**

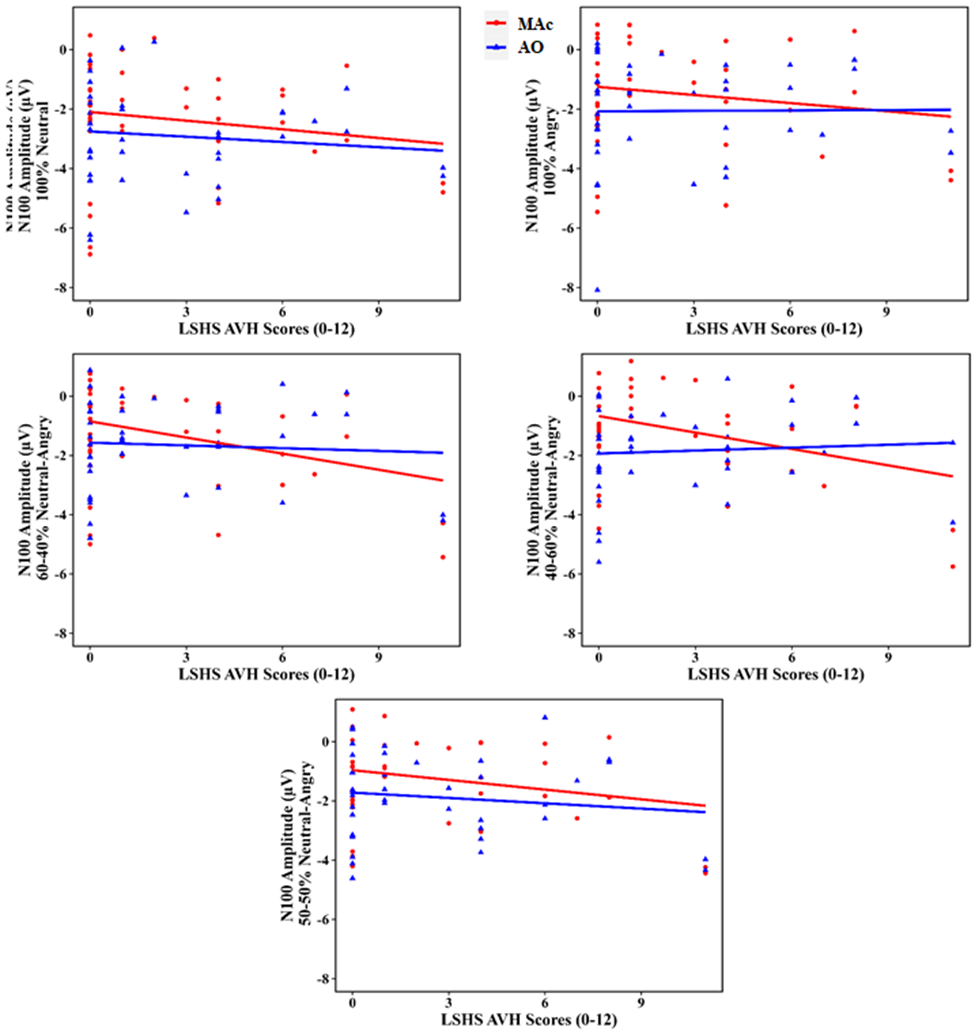

**Supplementary figure 6:** Scatter plots depicting the change in P200 amplitudes as a function of HP based on LSHS AVH scores for each stimulus type. Abbreviations: AO = Auditory Only; MAc = Motor Auditory corrected.

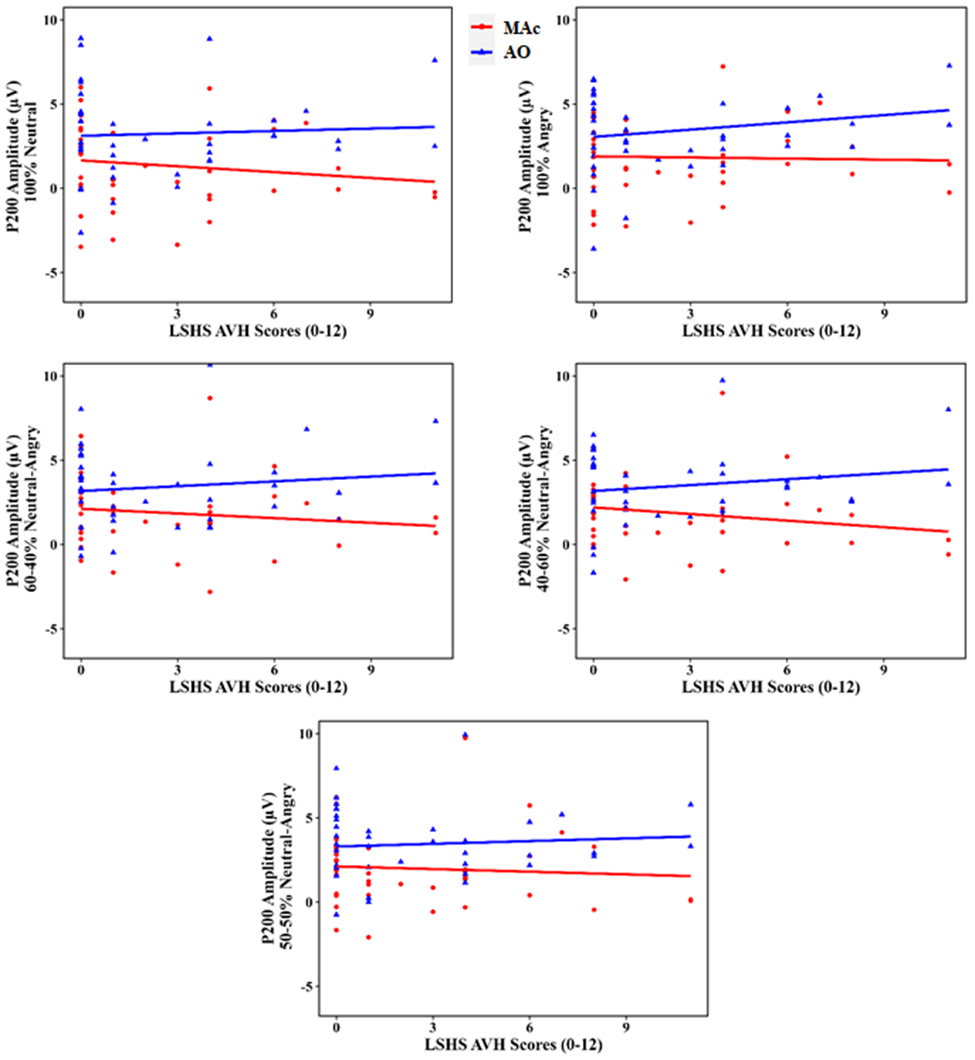

**Supplementary figure 7:** Scatter plots depicting the change in N200 amplitudes as a function of HP based on LSHS AVH scores for each stimulus type. Abbreviations: AO = Auditory Only; MAc = Motor Auditory corrected.

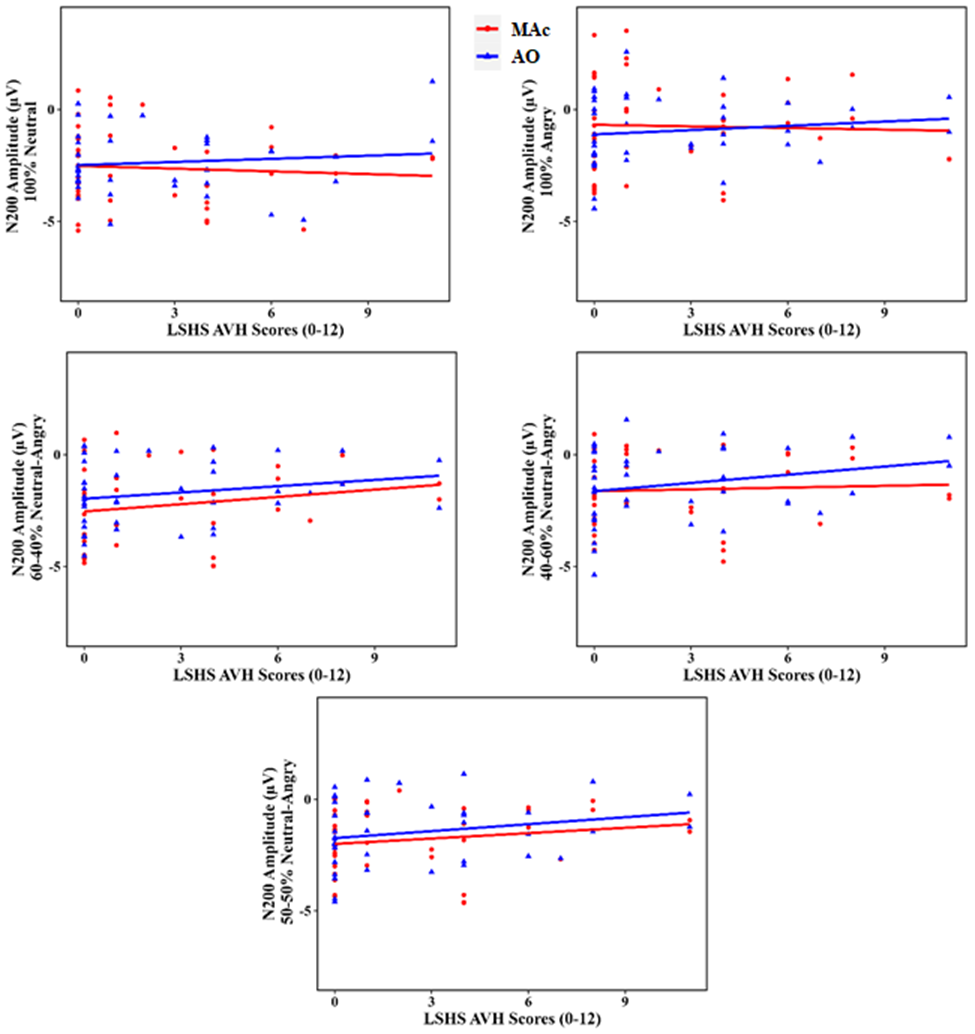
